# The Influence of Pesticide Use on Amphibian Chytrid Fungal Infections Varies with Host Life Stage

**DOI:** 10.1101/165779

**Authors:** Samantha L. Rumschlag, Jason R. Rohr

## Abstract

**Aim:** Pesticides are widespread and may alter host-pathogen interactions, ultimately influencing pathogen distributions across landscapes. Previous laboratory research supports two hypotheses regarding the effects of pesticides on interactions between amphibians and the predominately aquatic fungal pathogen *Batrachochytrium dendrobatidis* (Bd): 1) pesticides can be directly toxic to Bd reducing infection risk of aquatic larval amphibians, and 2) exposure to pesticides at formative stages of amphibian development can have long-term consequences on defenses, increasing disease risk after metamorphosis. It remains equivocal whether these laboratory patterns are consistent across amphibian species and occur in the field across broad spatial scales. The aim of this research is to address this research gap on the impact of pesticides on Bd distributions.

**Location:** Contiguous United States.

**Time Period:** 1998-2009.

**Major Taxa Studied:** Amphibian hosts and Bd.

**Methods:** Our data included 3,946 individuals evaluated for Bd infection across 49 amphibian species, at 126 locations, which resulted in 199 estimates of Bd prevalence in populations. We used species distribution models and multimodel inference to assess the influence of 1) total pesticide use, 2) pesticide use by type (herbicide, insecticide, fungicide), and 3) the most commonly used pesticide compounds on Bd infection prevalence in amphibian populations across life stages, controlling for several factors previously documented to affect Bd's distribution.

**Results:** Consistent with laboratory findings, our results indicate 36 that exposure to multiple herbicide compounds is associated with lowered infection risk in the aquatic larval stage but higher risk in the terrestrial post-metamorphic stage.

**Main Conclusions:** Our study highlights the complex nature of the effects that pesticides can have on disease distributions and suggests that pesticides should be strongly considered at broad scales and across host species, especially in environments in which exposure is widespread. Accurate predictions of disease distributions may lead to more effective management strategies to limit disease spread.

## INTRODUCTION

The emergence of infectious diseases threatens public health, global economies, and wildlife populations (Binder, 1999; Daszak *et al*., 2000; Morens *et al*., 2004). Therefore, understanding factors that determine distributions of infectious diseases is critical if we are to design effective management strategies to limit disease spread. Anthropogenic activities are predicted to be major determinants of infectious disease distributions (Daszak *et al*., 2001; Jones *et al*., 2008). While mounting evidence suggests that changes to climate and land use type can influence distributions of disease (Lafferty, 2009; Rohr *et al*., 2011; Martin & Boruta, 2013), the influence of chemical contaminants on disease distributions remains relatively undetermined (Lawler *et al*., 2006). For wildlife populations in freshwater ecosystems, chemical contaminants, including pesticides, are a widespread abiotic factor that might influence the distributions of disease occurrence by affecting host-pathogen interactions. Pesticides can have simultaneous positive and negative effects on parasite transmission; the net effect of these factors determines the influence of pesticides on disease risk in wildlife populations (Rohr et al., 2008a). For instance, pesticides can increase the incidence of pathogen infection (Christin et al., 2003; Pettis et al., 2012) via the disruption of host immune systems (Blakley et al., 1999; Rohr et al., 2008b). Alternatively, exposure to pesticides can also decrease pathogen viability via direct negative effects of pesticides on pathogen survival and reproduction (Lafferty & Kuris, 1999; Morley et al., 2003), pointing to the complex nature of effects of pesticides on host-pathogen interactions.

For hosts with complex life cycles, host life stage could determine the net effects of pesticides on disease risk if the relative balance between the effects of pesticides on host susceptibility and pathogen viability changes throughout the development of the host. If the net effect of pesticides changes with host life stage, we might expect pesticides to be negatively associated with infections for stages in which the negative effects of pesticides are greater on pathogen viability compared to host immunity. Alternatively, we might expect a positive association between pesticides and infection prevalence for host stages in which the negative impact of pesticides is greater on host immunity compared to pathogen viability.

Understanding the influence of pesticides on disease dynamics in amphibian populations at broad spatial scales is markedly important because amphibians are facing global declines that are caused, in part, by a fungal pathogen, *Batrachochytrium dendrobatidis* (hereafter, Bd) that causes the disease chytridiomycosis. Bd has been linked to population declines, mass mortality, and species extinctions in hosts (Lips *et al*., 2006; Skerratt *et al*., 2007), and its effects on hosts can be altered by environmental conditions, including pesticide exposure, that may influence pathogen viability or host immune response (Gaietto et al. 2014, Wise et al. 2014). Although the presence of Bd in North America dates back to 1888 (Talley *et al*., 2015), we lack an understanding of variation in host susceptibility across environmental gradients and the role pesticides might play in mediating the occurrence of Bd.

Experimental evidence suggests that pesticide exposure during critical developmental windows in early-life can have effects on the immune system in adult stages (Rohr & Palmer, 2005, 2013; Rohr *et al*., 2006) and that there are differential effects of pesticide exposure on amphibian-Bd interactions over aquatic larval and terrestrial post-metamorphic life stages. For instance, in the aquatic larval life stage of amphibians, pesticides can have direct negative effects on Bd, which results in an overall decreased risk of disease for aquatic larvae. In fact, Bd growth on infected tadpoles can be reduced by pesticide exposure (McMahon *et al*., 2013) and can even result in clearance of Bd from the host (Hanlon *et al*., 2012). These negative effects of pesticides on Bd are likely driven by reduced Bd growth and production of Bd zoospores, the aquatic infective stage of the pathogen (Hanlon *et al*., 2012; McMahon *et al*., 2013). Bd infects keratinized cells in amphibians, which occur only in the mouthparts of tadpoles (Voyles *et al*., 2011), suggesting that susceptibility to Bd infection is low in this early-life stage. As tadpoles metamorphose into the terrestrial life stage, the incidence of keratinized cells increases as the epidermis develops and Bd infection can move from the mouthparts of the tadpole to the entire surface of the body (McMahon & Rohr, 2015), suggesting that susceptibility to infection and disease development increases in the terrestrial host life stage (Rachowicz & Vredenburg, 2004). In the terrestrial post-metamorphic life stage, pesticide exposure during early-life is associated with increased Bd-induced mortality, which may be driven by disruption of the immune system. For example, early-life pesticide exposures can lead to increased Bd-induced mortality of terrestrial hosts, which is caused by reduced tolerance to infection; this finding points to a cost of pesticide exposure that could be induced by disruption of the immune system (Rohr *et al*., 2013). While these experimental studies support differential effects of pesticide exposure on amphibian-Bd interactions over aquatic larval and terrestrial post-metamorphic life stages, it remains equivocal whether these laboratory patterns are consistent across amphibian species and occur in natural populations at broad spatial scales.

The objective of the current study is to evaluate the influence of pesticide use on Bd infection prevalence in amphibian populations across the United States. We used publically available data including 199 field observations of Bd infection prevalence and corresponding estimates of pesticide use at the county level. We used species distribution models and multimodel inference approaches to assess the influence of 1) total pesticide use, 2) pesticide use by type (herbicide use, insecticide use, fungicide use) and 3) the most commonly used pesticide compounds within type on Bd infection prevalence in amphibian populations across life stages controlling for the influence of environmental (vegetation, precipitation, temperature) and biotic (host family) factors. Based on the experimental evidence reviewed previously concerning persistent and life-stage dependent effects of pesticides on Bd infection risk in amphibians, we predicted that in the aquatic larval stage of amphibians, Bd prevalence would be negatively associated with pesticide use, and in the terrestrial post-metamorphic stage, Bd prevalence would be positively associated with pesticide use.

## METHODS

### Response and Predictor Variables

We obtained a spatially explicit dataset of amphibian populations surveyed for Bd infection from Bd Maps (www.bd-maps.net) in 2013. Bd survey sites were included in our analyses if five or more individuals were surveyed at a given site between 1992 and 2012, life stage information of amphibians was provided, and survey sites were located in the contiguous USA. The resulting dataset comprised 3,946 individuals evaluated for Bd infection, across 49 species, at 126 unique locations, which resulted in 199 observations of Bd infection prevalence at the population level. Bd infection prevalence at each site, which served as the response variable in all of our statistical models, was arcsine-square-root transformed for each analysis.

To conservatively estimate pesticide use, we used USA county-level low pesticide use (as opposed to high) estimates from 1992 to 2012 obtained from the Estimated Annual Agricultural Pesticide Use dataset provided by the Pesticide National Synthesis Project of the National Water-Quality Assessment (NAWQA) Program (United States Geological Survey) (https://water.usgs.gov/nawqa/pnsp/usage/maps/county-level/). Preliminary analyses showed the effects of high pesticide use estimates were indistinguishable from low use estimates (data not shown). We classified the pesticide compounds as herbicide, insecticide, or fungicide using the primary use type classifications provided by Pesticide Action Network (PAN) Pesticide Database (http://www.pesticideinfo.org/). We included plant growth regulators and defoliants as herbicides and insect growth regulators as insecticides. For a given site in a given county, we summed low use estimates across pesticide types at the county level. In statistical models, pesticide usages were transformed using the natural logarithm. We excluded mineral and biologic fungicides (e.g. bacteria) because we were interested in non-target effects of synthetic fungicides on host responses to Bd that might influence infection prevalence. To estimate local vegetative habitat, we used a seven-day composite from June 14 to 20 of 2002 of Normalized Difference Index (NDVI) data from eMODIS (Earth Resources Observation and Science Center, Moderate Resolution Imaging Spectroradiometer) made available by the United States Geological Survey (https://earthexplorer.usgs.gov/). Data for the following abiotic factors were downloaded from WorldClim (http://www.worldclim.org/): 30-y means (1960-1990) of annual total precipitation, precipitation of the wettest month, precipitation of the driest month, annual mean temperature, mean diurnal temperature range, maximum temperature of the warmest month, and minimum temperature of the coldest month. NDVI, precipitation, and temperature measures were extracted at survey site locations using ArcMap 10.4. We reduced our three precipitation measures and four temperature measures into a single precipitation measure and a single temperature measure using principal component analyses by extracting the first axis for precipitation measures (98.9% of the total variation, hereafter PC1Precip) and for temperature measures (83.8% of the total variation, hereafter PC1Temp).

### Generalized Least Squares Models

Generalized least squares (GLS) multiple regression models were fit using the gls function (nlme package) with full maximum likelihood fit and an exponential spatial correlation structure. In all models, observations were weighted based on the number of individuals surveyed at a site. We constructed three sets of models to evaluate the influence of 1) total pesticide use, 2) pesticide use by type (herbicide use, insecticide use, fungicide use), and 3) the most commonly used herbicide compounds on Bd infection prevalence across host life stages. Models for total pesticide use and pesticide use by type included the following covariates to control for the effects of biotic and environmental factors: host family, NDVI, PC1Precip. and PC1Temp. To simplify models of pesticide compounds, we included host family and PC1Temp. as the only biotic and environmental covariates, as these covariates were relatively important in models of total pesticide use and pesticide use by type (relative importance score > 0.6, Figure 1).

**Fig. 1.**
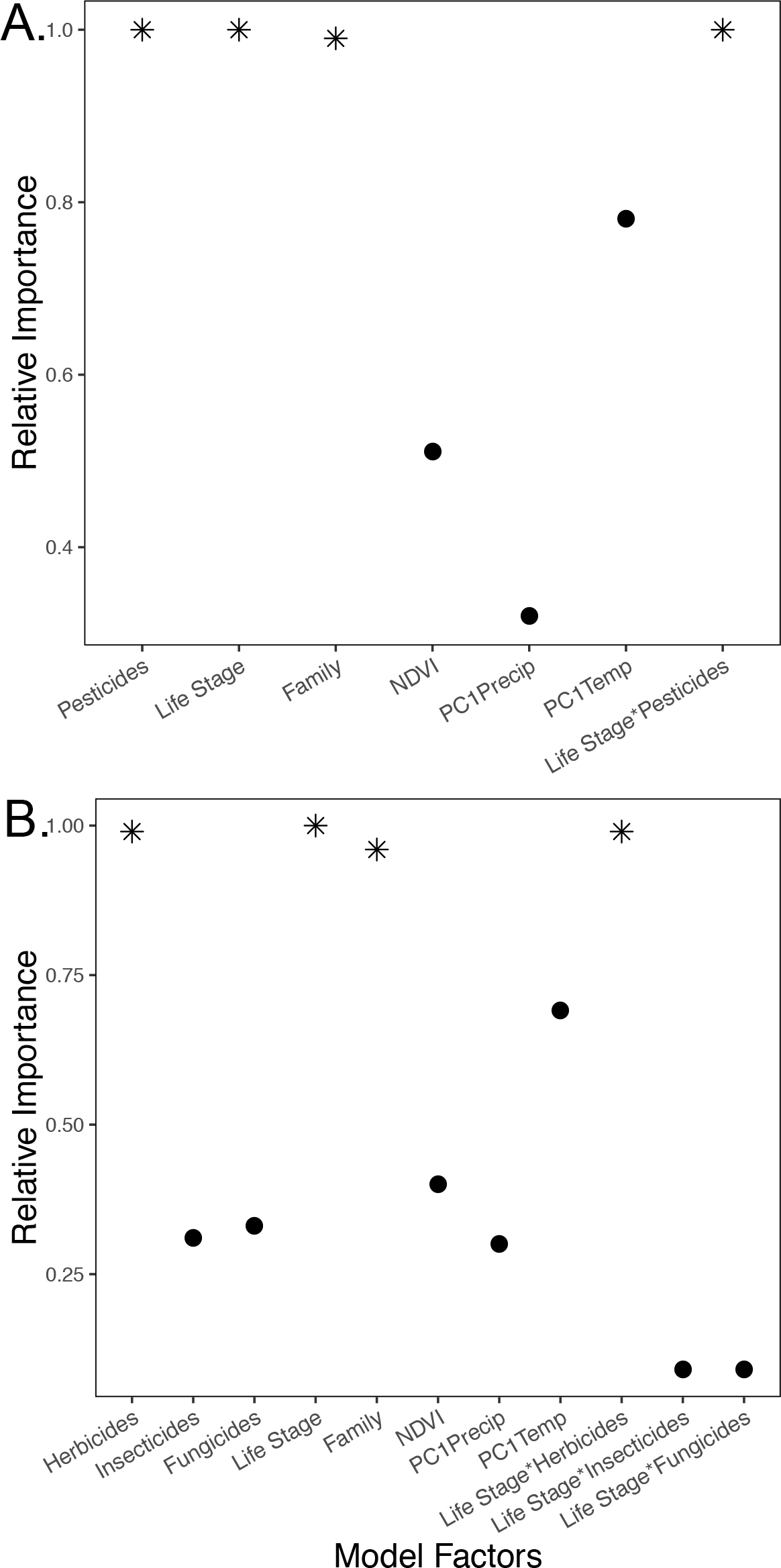
Relative importance of factors included in model comparisons evaluating the influence of A) total pesticide use across host life stages and B) herbicide, insecticide, and fungicide use across host life stages and model covariates (family, NDVI, precipitation, and temperature). Stars indicate significance of the factor (*p* < 0.05) from model averaging.

To evaluate the influence of total pesticide use across life stages of amphibian hosts on Bd infection prevalence, we constructed a model with predictor variables including: total pesticide use, host life stage, the interaction of life stage and pesticides, and all covariates. To evaluate which type of pesticide drove the effects of total pesticide use on Bd infection prevalence across life stages, we constructed a model with predictor variables including: herbicide use, insecticide use, fungicide use, host life stage, the two way interactions of host life stage with herbicide, insecticide, or fungicide use, and all covariates. When controlling for the effects of insecticide and fungicide use, herbicide use was the best predictor of Bd infection prevalence across host life stages. To determine which commonly used herbicide compounds drove the effects of herbicide use, we first gathered the use estimates for the top five most-used herbicide types in our dataset and constructed five models that included the following predictor variables: herbicide compound (glyphosate, atrazine, metolachlor-s, ethephon, or sodium chlorate), herbicide use minus the compound of focus, insecticide use, fungicide use, host life stage, the interaction of host life stage and herbicide compound of focus, host family, and PC1Temp. For use estimates at sites in which a compound estimate was not given, we assumed no use of that compound for the given site.

### Multimodel Inference and Comparisons of Goodness of Fit

To avoid relying on a single model to draw conclusions about the importance of predictors on prevalence, we used multimodel inference (MuMin package), which fits models using combinations of predictors and ranks models by second-order Akaike Information Criteria corrected for small sample sizes (AICc) (dredge function), for models including total pesticide use and pesticide use by type. AICc, ΔAICc, and Akaike weights for each candidate model were calculated. To compare the influence of model factors across all candidate models, Akaike weights for each factor were summed across models to determine relative importance scores (Burnham & Anderson, 2002). P-values were calculated from full model-averaged parameter estimates with statistical shrinkage. Nagelkerke pseudo R^2^ values were calculated to assess goodness-of-fit of the top performing models (with a ΔAICc equal to zero). To visualize the effect of significant predictors on prevalence, we provide partial regression plots from these top performing models including total pesticide use and use by pesticide type.

To determine the relative contribution of each of the top five herbicide compounds to the patterns of total herbicide use on Bd prevalence across host life stages, we used a log-likelihood ratio test to compare the goodness-of-fit of two models: one that included the interaction between the focal herbicide compound and host life stage and one that did not. To visualize the influence of herbicide compounds on infection prevalence, we provide partial regression plots for each herbicide compound and the sum of the most used herbicide compounds across host life stages controlling for covariates.

## RESULTS

### The Influence of Total Pesticides

Total pesticide use, life stage, family, and the interaction of pesticide use and life stage influenced Bd infection prevalence in amphibian populations significantly when controlling for covariates (Table 1, Fig. 1A, Nagelkerke pseudo R^2^ = 0.39). The relative importance scores for pesticide use, life stage, and the interaction of pesticide use and life stage were greater than all covariates, including host family, NDVI, precipitation, and temperature (Fig. 1A). In the best-fitting model controlling for covariates, the impact of pesticides depended on life stage (Table 1, Fig. 2A). For the aquatic larval life stage of the hosts, Bd infection prevalence decreased with increasing pesticide use, but for the terrestrial post metamorphic life stage of hosts, Bd infection prevalence increased with pesticide use (Fig. 2A).

**Table 1.** Model averaged coefficients, standard error (SE), z-statistics and associated *p*-values with statistical shrinkage from multimodel inference analyses predicting the effects of total pesticide use and pesticide use by type on Bd prevalence in amphibian populations.

| Variable | Coefficient | SE | z | *p* |
| --- | --- | --- | --- | --- |
| **Total Pesticide Use** |  |  |  |  |
| (Intercept) | 1.0968 | 0.2776 | 3.93 | **<0.001** |
| Pesticides | -0.0811 | 0.0238 | 3.38 | **0.001** |
| Life Stage (Terrestrial) | -1.2164 | 0.3135 | 3.86 | **<0.001** |
| Family (Hylidae) | -0.1427 | 0.0931 | 1.52 | 0.128 |
| Family (Plethodontidae) | -0.2146 | 0.0947 | 2.25 | **0.024** |
| Family (Ranidae) | 0.0145 | 0.0808 | 0.18 | 0.858 |
| Family (Salamandridae) | 0.1392 | 0.1573 | 0.88 | 0.379 |
| NDVI | -0.0811 | 0.1105 | 0.73 | 0.464 |
| PC1Precip. | 0.0016 | 0.0051 | 0.30 | 0.761 |
| PC1Temp. | -0.0592 | 0.0453 | 1.30 | 0.193 |
| Pesticides*Life Stage (Terrestrial) | 0.1396 | 0.0293 | 4.74 | **<0.001** |
| **Type of Pesticide Use** |  |  |  |  |
| (Intercept) | 0.9807 | 0.2510 | 3.88 | **<0.001** |
| Herbicides | -0.0704 | 0.0245 | 2.86 | **0.004** |
| Insecticides | -0.0007 | 0.0173 | 0.04 | 0.970 |
| Fungicides | -0.0033 | 0.0127 | 0.26 | 0.797 |
| Life Stage (Terrestrial) | -1.1068 | 0.3041 | 3.62 | **<0.001** |
| Family (Hylidae) | -0.1338 | 0.0951 | 1.40 | 0.162 |
| Family (Plethodontidae) | -0.2262 | 0.1023 | 2.20 | **0.028** |
| Family (Ranidae) | -0.0070 | 0.0813 | 0.09 | 0.932 |
| Family (Salamandridae) | 0.0937 | 0.1565 | 0.60 | 0.552 |
| NDVI | -0.0504 | 0.0931 | 0.54 | 0.590 |
| PC1Precip. | 0.0011 | 0.0047 | 0.24 | 0.813 |
| PC1Temp. | -0.0469 | 0.0442 | 1.06 | 0.290 |
| Herbicides*Life Stage(Terrestrial) | 0.1315 | 0.0332 | 3.94 | **<0.001** |
| Insecticides*Life Stage (Terrestrial) | 0.0019 | 0.0172 | 0.11 | 0.912 |
| Fungicides*Life Stage (Terrestrial) | 0.0017 | 0.0109 | 0.16 | 0.875 |

**Fig. 2.**
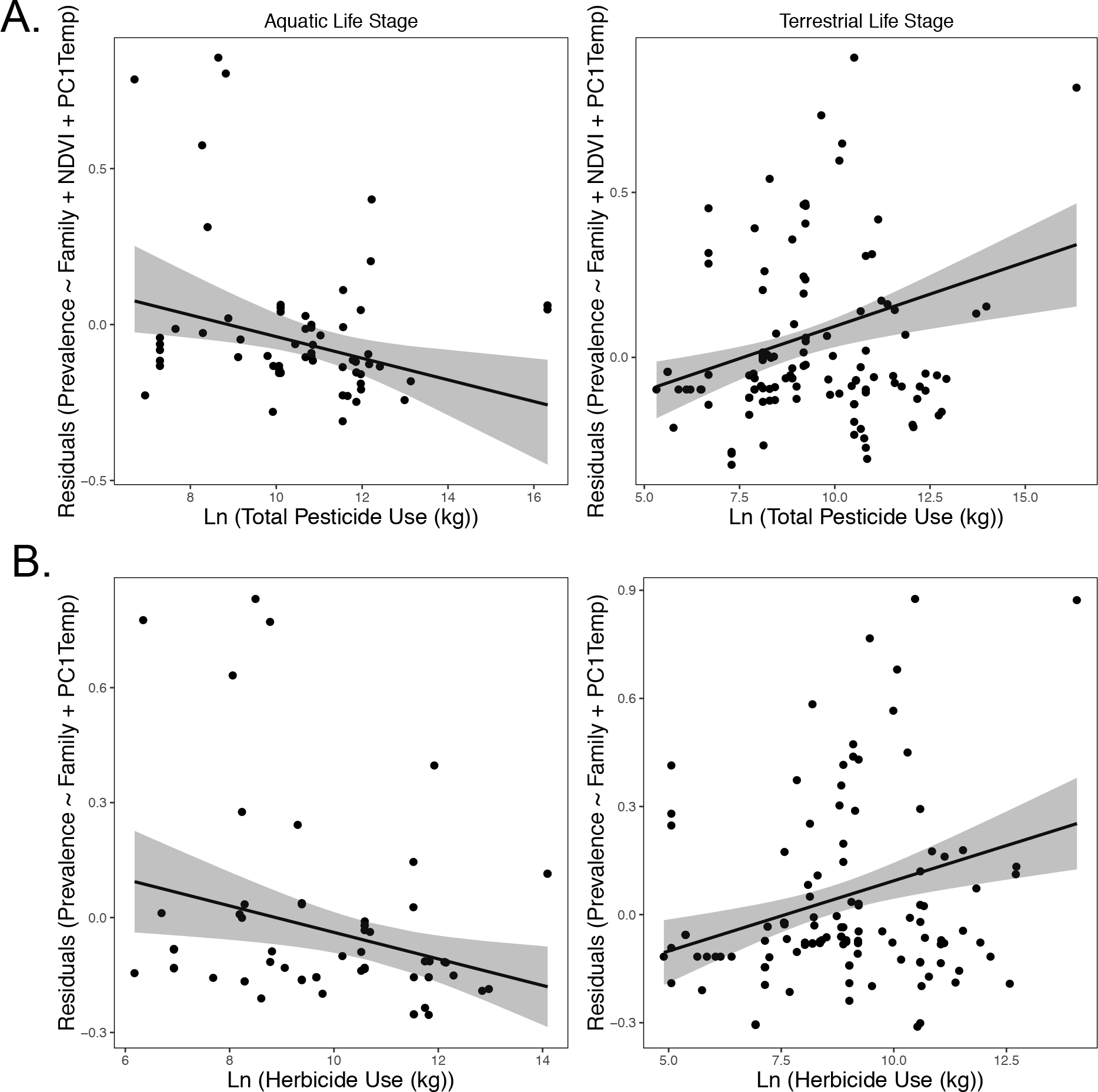
Partial regressions showing A) the influence of total pesticide use on *Batrachochytrium dendrobatidis* infection prevalence in amphibian populations across aquatic and terrestrial amphibian life stages controlling for effects of host family, NDVI, and temperature and B) the influence of herbicide use on *Batrachochytrium dendrobatidis* infection prevalence in amphibian populations across aquatic and terrestrial life stages controlling for effects of host family and temperature. Models shown are the models with a ΔAICc equal to zero from model comparisons. Prevalence has not been transformed. Gray bands represent 95% confidence intervals.

**Table 2.** Model comparison results for models with ΔAICc values less than 2 testing predictors including total pesticides and types of pesticides.

| Models Including Total Pesticide Use | AICc | $\Delta\text{AICc}$ | w |
| --- | --- | --- | --- |
| Family + Life Stage + Pesticides + NDVI + PC1Temp + Pesticides*Life Stage | 192.21 | 0 | 0.28 |
| Family + Life Stage + Pesticides + PC1Temp + Pesticides*Life Stage | 192.48 | 0.27 | 0.24 |
| Family + Life Stage + Pesticides + NDVI + PC1Temp. + PC1Precip. +<br>Pesticides*Life Stage | 193.47 | 1.26 | 0.15 |
| Family + Life Stage + Pesticides + PC1Temp. + PC1Precip. + Pesticides*Life Stage | 194.05 | 1.84 | 0.11 |
| Models Including Types of Pesticides |  |  |  |
| Family + Life Stage + Herbicides + PC1Temp. + Herbicides*Life Stage | 193.77 | 0 | 0.12 |
| Family + Life Stage + Herbicides + NDVI + PC1Temp. + Herbicides*Life Stage | 194.34 | 0.57 | 0.09 |
| Family + Life Stage + Herbicides + Herbicides*Life Stage | 194.92 | 1.16 | 0.07 |
| Family + Life Stage + Herbicides + PC1Precip. + PC1Temp. + Herbicides*Life<br>Stage | 195.45 | 1.68 | 0.05 |
| Family + Life Stage + Herbicides + Fungicides + PC1Temp. + Herbicides*Life Stage | 195.67 | 1.91 | 0.05 |
| Family + Life Stage + Herbicides + NDVI + PC1Precip. + PC1Temp. +<br>Herbicides*Life Stage | 195.74 | 1.98 | 0.04 |

### The Influence of Pesticide Use by Type

While pesticide uses by type were positively correlated (herbicide use vs. insecticide use: Pearson’s correlation coefficient = 0.77, herbicide use vs. fungicide use = 0.59, insecticide use vs. fungicide use = 0.82), the influence of total pesticide use on Bd infection prevalence across host life stages seemed to be driven by herbicide use in comparison with insecticide and fungicide use (Table 1). Herbicide use, life stage, family, and the interaction of herbicide use and life stage were significant predictors of prevalence, controlling for insecticide use, fungicide use, the interaction of insecticide and fungicide uses with life stage, and all other covariates (Table 1). The relative importance scores for herbicide use, life stage, and the interaction of herbicide use and life stage were greater than all other factors in the model, including family, insecticide use, fungicide use, and the interaction of insecticide or fungicide use with life stage (Fig. 1B). Similar to the effect of total pesticide use, herbicide use was associated with decreased infection prevalence in the aquatic larval life stage and increased infection prevalence in the terrestrial post-metamorphic life stages (Fig. 2B, Nagelkerke pseudo R^2^ = 0.37).

### The Influence of Herbicide Compounds

The five most commonly used herbicides in the dataset include glyphosate (34% of total herbicide use based on weight), atrazine (10%), metolachlor-s (5%), ethephon (5%), and sodium chlorate (5%). Including the interaction between focal herbicide compound and life stage improved the goodness of fit compared to the same model without this interactions for all five herbicides (glyphosate [log-likelihood ratio = 17.55, *p*<0.001], atrazine [log-likelihood ratio = 8.97, *p* = 0.003], metolachlor-s [log-likelihood ratio = 8.58, *p* = 0.003], ethephon [log-likelihood ratio = 4.89, *p* = 0.027], sodium chlorate [log-likelihood ratio = 6.09, *p* = 0.01]). Similar to the effect of total pesticides and herbicides, glyphosate use was negatively associated with Bd prevalence in the aquatic stage and positively associated in the terrestrial stage (Fig. 3A). Both atrazine and metolachlor-s use were negatively associated with Bd prevalence in the larval stage but did not appear to be associated with infections in the terrestrial stage (Fig. 3B,C). In contrast, ethephon and sodium chlorate use did not have a strong influence on Bd prevalence in the aquatic larval stage, but were positively associated with Bd prevalence in the terrestrial post-metamorphic stage (Fig. 3D,E). The influence of the sum of the top five most-used herbicide compounds matches closely with the pattern of overall herbicide use on Bd prevalence (Fig. 3F).

**Fig. 3.**
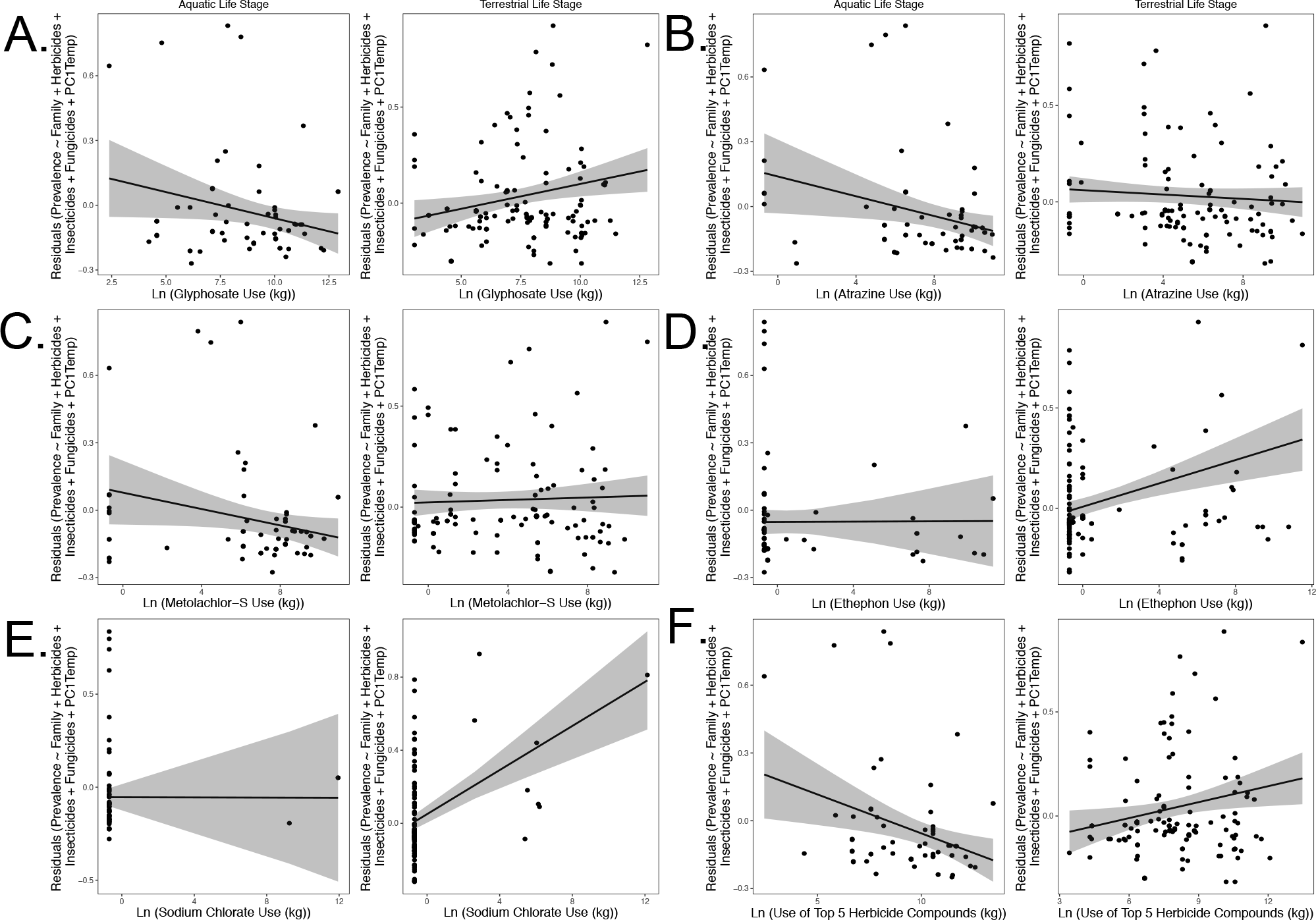
Partial regressions showing the influence of A) glyphosate use, B) atrazine use, C) metolachlor-s use, D) ethephon use, E) sodium chlorate use, and F) the sum of these top five herbicide compounds across life stages on Bd infection prevalence in amphibian populations controlling for the effects of family, herbicide use (minus the compound or group of focus), insecticide use, fungicide use, and temperature. Prevalence has not been transformed. Gray bands represent 95% confidence intervals.

## DISCUSSION

Pesticides represent a major ecological disturbance to communities in aquatic environments and can shape distributions of organisms across landscapes (Liess *et al*., 2005; Schäfer *et al*., 2007; Beketov *et al*., 2013). Our study provides evidence of the influence of pesticides on infectious disease dynamics of wildlife at broad spatial scales, which is consistent with the body of experimental research on pesticides and amphibian-Bd interactions. We show a negative relationship between pesticide use and Bd infection prevalence in the aquatic larval life stage and a positive relationship between pesticide use and Bd infection prevalence in the terrestrial post-metamorphic life stage. Our analyses suggest that the combined influence of the most commonly used herbicides are the primary determinants of these differential effects of pesticides on infection prevalence across host life stages.

Hosts and pathogens in freshwater systems are likely exposed to pesticides in the aquatic environment, where the presence of contaminants, including pesticides, is common because of aerial deposition and agricultural runoff (Gilliom & Hamilton, 2006). In the amphibian-Bd system, when pesticides are present, the balance between hosts and pathogens in the aquatic environment is likely tipped in favor of hosts because of direct negative effects of pesticides on pathogen viability, which explains the mechanism for the observed negative association between pesticide use and infection prevalence in the aquatic larval stage of hosts in the current study.

Several pesticides, including atrazine—the second most used herbicide compound in the United States and in our dataset, have been shown to have direct negative effects on Bd growth, survival (Hanlon & Parris, 2012; McMahon *et al*., 2013), and production of zoospores, the aquatic infective stage of Bd (Hanlon & Parris, 2012).

However, as hosts develop, they can suffer delayed negative effects of early-life pesticide exposure well into adulthood, increasing their overall risk of disease development in the terrestrial post-metamorphic life stage. Pesticide exposure can have delayed effects on organisms (Rohr & Palmer, 2005; Jones *et al*., 2009) and can disrupt host-pathogen interactions leading to an increase in infectious disease risk (Rohr *et al*., 2006; Rohr & McCoy, 2010; Wise *et al*., 2014). We propose that the observed positive effect of pesticide use, mainly driven by herbicide use, on Bd infection prevalence is consistent with the body of research showing persistent negative effects of early-life exposures to pesticides on infectious disease risk; for instance, post-metamorphic amphibians can suffer increased mortality to Bd as a result of early-life exposure to atrazine caused by reduced tolerance to infection, suggesting a long-term cost of pesticide exposure (Rohr *et al*., 2013).

Interestingly, herbicides, as opposed to insecticides or fungicides, were most correlated with the observed patterns between pesticide use and infection prevalence. We suspect the power to detect an effect of herbicides is greater than that for insecticides or fungicides because herbicides are used in greater amounts in the United States (Grube *et al*., 2011), increasing the likelihood of exposure in natural systems. In a given year, herbicides are used more than five times as much as insecticides or fungicides as measured by mass of active ingredient (Grube *et al*., 2011). Herbicide exposure of natural host-pathogen populations is therefore more likely to occur in comparison with exposure to insecticides or fungicides.

Our results support that the combined effects of the most commonly used herbicides together drive the observed patterns of total herbicide use on infection prevalence. The association between individual herbicide compounds and infection prevalence across life stage either closely matched the overall pattern of total herbicide use (e.g. glyphosate) or showed a similar pattern to the influence of total herbicide use in at least one of the host life stages (e.g. atrazine, metolachlor-s, ethephon, sodium chlorate). For instance, atrazine and metolachlor-s have negative effects on infection prevalence in the aquatic larval life stage but no strong effect in the terrestrial post-metamorphic life stage, and ethephon and sodium chlorate have positive effects on infection prevalence in the terrestrial post-metamorphic life stage but no strong effect in the aquatic larval life stage. The top five most commonly used herbicides in our dataset comprised about 59% of the total use of herbicides, so when we examine the influence of the sum of these herbicide compounds on infection prevalence across host life stages (Fig. 3F), it unsurprisingly closely matched the patterns for total herbicide use (Fig. 2B).

Our results suggest that managers could favor certain herbicide compounds over others if their goal is to limit increasing susceptibility to Bd infection in the terrestrial post-metamorphic stage. An understanding of non-target effects, including the role of potential herbicides on amphibian host fitness, would be needed to develop an integrated pest management solution.

While our study evaluates the potential influence of pesticides on amphibian host resistance, via infection prevalence, we have not evaluated fitness consequences of pesticides on hosts exposed to parasites, which may occur through the physiological mechanisms of resistance or tolerance. For instance, herbicide exposure of Bd-infected or -exposed amphibian hosts may result in increased host mortality.

While we support that the combined uses of the most common herbicide compounds drive the influence of pesticide use on Bd infection prevalence in amphibians, we do not suggest that insecticides and fungicides do not contribute to this pattern. Instead, we highlight that herbicide, insecticide, and fungicide use are positively correlated at the county level across the United States. Since pesticide use estimates are derived at least in part from land use data (Thelin & Stone, 2013), counties in which herbicide use is high are likely counties with increased agricultural land use, so these counties also have high insecticide and fungicide use. The influence of the most abundant pesticide type, namely herbicides, gives rise to the model that best predicts infection prevalence. Even though our models control for the use of other pesticide types when testing for a focal pesticide type, because of the positive correlation among the pesticide use types, we are hesitant to disregard a potential influence of insecticides and fungicides on Bd distributions.

Pesticides can be a major driver of communities in freshwater ecosystems (McMahon et al. 2012). Consistent with the body of experimental evidence in this system, our research illustrates how pesticides can shape distributions of infectious pathogens over broad spatial scales via effects that vary over the life span of a host, which highlights the complex nature of the impact of contaminants on natural systems. With their impacts on pathogen viability and host immunity, the effects of pesticides on infectious disease distributions should be given more attention particularly at broad scales and across host species. Accurate predictions of disease distributions may lead to the most effective management strategies to limit the spread of diseases to vulnerable populations.

## ACKNOWLEDGMENTS

— Thank you to M. Venesky for providing the Bd dataset and to K. Ronnenberg and D. Olson for information regarding updates to the Bd dataset. We are grateful for the thoughtful advice of S. Koerner, J. Cohen, and M. Mahon on data management and analyses and to the many scientists who freely shared their data by contributing to the Bd-maps database. This manuscript was improved by the thoughtful feedback of the Rohr lab.

## BIOSKETCH

Samantha Rumschlag (samantharumschlag.weebly.com) is currently a postdoctoral scholar in Jason Rohr’s lab at the University of South Florida. Her research combines experimental manipulations with broad scale analyses to examine the influence of anthropogenic changes on wildlife populations and infectious diseases. The goal of her research is to inform wildlife management concerning current environmental issues to increase the likelihood of sustainable coexistence between humans and wildlife.

## DATA ACCESSIBILITY

The derived dataset used for these analyses will be made publically available via Dryad upon manuscript acceptance.

